# Automatic Sleep Stage Classification with Optimized Selection of EEG Channels

**DOI:** 10.1101/2022.06.14.496176

**Authors:** Håkon Stenwig, Andres Soler, Junya Furuki, Yoko Suzuki, Takashi Abe, Marta Molinas

**Affiliations:** Department of Engineering Cybernetics, Norwegian University of Science and Technology, Trondheim, Norway; International Institute for Integrative Sleep Medicine (WPI-IIIS), University of Tsukuba, Tsukuba, Japan

**Keywords:** Polysomnography, NSGA, Machine Learning, sleep scoring, EEG

## Abstract

Visual inspection of Polysomnography (PSG) recordings by sleep experts based on established guidelines has been the gold standard in sleep stage classification. This approach is expensive, time consuming and mostly limited to experimental research and clinical cases of major sleep disorders. Various automatic approaches to sleep scoring have been emerging in the past years and are opening the way to a quick computational assessment of sleep architecture that may find its way to the clinics. With the hope to make sleep scoring a fully automated process in the clinics, we report here an ensemble algorithm that aims at not only predicting sleep stages but of doing so with an optimized minimal number of EEG channels. For that, we combine a genetic algorithm based optimization with a classification framework that minimizes the number of channels used by the machine learning algorithm to quantify sleep stages. This resulted in a scoring with an F1 score of 0.793 for the fully automatic model and 0.806 for the model trained on 10 percent of the unseen subject, both with only 3 EEG channels. The ensemble algorithm is based on a combination of extremely randomized trees and MiniRocket classifiers. The algorithm was trained, validated and tested on night sleep PSG data collected from 7 subjects. The novelty of our approach lies on the use of the minimum information needed for automated sleep scoring, based on a systematic search that concurrently selects the optimal-minimum number of EEG channels and the best performing features for the machine learning classifier. The optimization framework presented in this work may enable new designs for sleep scoring devices suited to studies in the comfort of the homes, easily and inexpensively and in this way facilitate experimental and clinical studies in large populations.

## I. Introduction

The quality of sleep is crucial for overall health of human beings and is becoming one of the top public health concerns. Altered sleep patterns affect people’s daily performance and several health issues are closely associated with poor sleep quality. Some neurological diseases, cardiovascular and metabolic disorders and weakened immune system have been associated with sleep-related disorders [1]–[3]. Early detection of sleep pattern alterations may prevent further evolution of these disorders. The first step in any sleep study is the annotations of the different sleep stages, which is typically performed by visual examination of PSG recordings as the gold standard of sleep assessment [4], [5]. PSG monitors brain activity (EEG), muscle activity (EMG) and eye movements (EOG) and it typically requires that the patients sleep overnight at the hospital or sleep laboratory while their signals are recorded.

The annotations of stages are then performed manually by well-trained human experts, a lengthy and tedious task that also generates considerable inter-rater variability [6]. Many different automatic scoring approaches based on machine learning and deep learning algorithms have been proposed over the years with reasonably good scoring accuracies [7]– [13], but as of today automatic scoring is not a widespread practice in the sleep clinics. Besides accuracy, the adoption of automatic sleep scoring depends among other things, on some practical aspects of the implementation such as the associated computational costs when using all the PSG channels, and the simplicity of the recording device [8], [14]. Most of these reduced channels were selected a-priori based on the experience of the experts. Many alternative methodologies based on reduced channels or EEG-only channels are discussed in the literature and they obtain reasonably high annotation accuracy of sleep stages [8]. However, a systematic selection of the optimal channels that contain the most relevant information to increase the classification accuracy, searching on the entire high density EEG space (e.g. 128ch), has not been investigated until now. Collecting the minimum information needed for automated sleep scoring may facilitate its adoption in clinics in the future. This work aims at providing such a methodology by combining optimal EEG channel selection with optimally selected features for the machine learning algorithms, to achieve reasonably high scoring accuracy with a minimal number of EEG channels. Selecting EEG channels according to their contribution to classification accuracy, will warrant the minimal information required for the task while significantly reducing computation and increasing the likelihood of real-time implementations. This optimization-based automated sleep scoring algorithm may also be used as a platform for designing new and simplified sleep scoring devices that can facilitate sleep studies at home while retaining high accuracy and by that accelerate experimental research and sleep studies across large populations.

## II. Materials and Methods

### A. Polysomnographic (PSG) Data

The sleep dataset used for training and testing was recorded at the Human Sleep Lab of the International Institute of Integrative Sleep Medicine. It is obtained through PSG recordings of 7 subjects, each with 136 channels, 128 EEG channels, 2 mastoid channels, 3 Electromyography (EMG) channels, and 3 Electrooculography (EOG) channels. The sampling rate for the dataset is 1024 Hz. The average age of the seven subjects is 22.4 ± 0.8 years, with the age range of 22-24, and the number of epochs totals to 6955. It includes 3 males, with average age of 22.0 ± 0.0 years and age range (22), and 4 females, with average age of 22.8 ± 1.0 years, and age range of 22-24. The EEG channels are located according to the biosemi128 configuration^1^. All datasets were scored by a sleep expert in 30 second epochs according to the AASM rules [4].

The distribution of the sleep stages for each subject and for the entire dataset can be seen in figure 1. All the subjects combined exhibit a fairly normal sleep stage distribution [15], apart from a slightly higher occurrence of N1 and wakefulness, but this is partly due to subjects 3 and 4 exhibiting a higher frequency of N1 and subject 4 having an unusual distribution, almost uniformly distributed among all the stages.

**Fig. 1.**
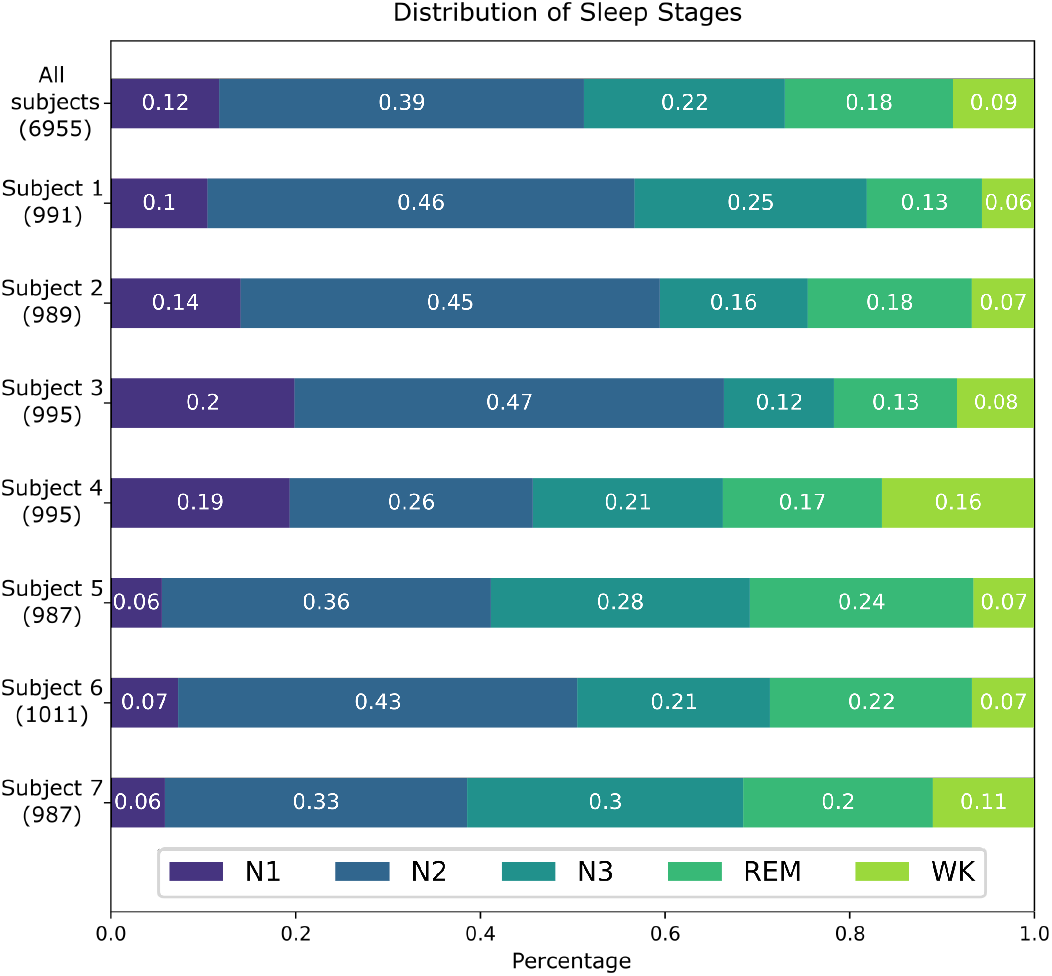
The distribution of sleep stages for each subject and the whole dataset. The number of epochs are in parentheses

The Montreal Archive of Sleep Studies (MASS) cohort was used for training the spindle detector (SS2 subset). The SS2 subset includes 19 subjects, average age of 23.6 ± 3.7 years, and age range of 18-33. It includes 8 males, average age of 24.3 *±* 4.2 years and age range of 19-33, and 11 females, average age of 23.2 *±* 3.5 years and age range of 18-30.

### B. Performance Evaluation

The performance of a classifier can be determined through different performance measures. Commonly, accuracy is atilized, which is the fraction of predictions the model got right. The accuracy as a performance measure is sensitive to class imbalance: If a subject had 50 percent of epochs belonging to the N2 class a classifier which only predicted N2 would achieve an accuracy of 50 percent. And because of the class imbalance for this classification problem other performance measures are necessary to provide a more nuanced picture.

Two other important measures are precision and recall. Precision is a measure of how many predictions are actually positive of all the positive predictions, while recall is a measure of how many of all the positive cases are predicted as positive. Precision and recall can be calculated for binary classification problems while for multiclass classification they can be calculated per class. These two metrics are generally competing metrics, predicting every epoch as one class will give this class a high recall score, but a low precision score. These two measures can be combined into the F1 score which is the harmonic mean of the precision and recall. Thus, a high F1 score will reflect both a few number of both false positives and false negatives.

#### 1) Training,Validation and testing

Cross-validation is a technique used to evaluate the performance of a machine learning model. It is commonly applied to predictive models, because it is easy to implement and generally it has a lower bias than other methods, such as a simple train and test split. The objective of cross-validation is to test the model’s ability to predict new data that was not used in estimating it, to identify problems like overfitting and selection bias. An extension of Cross-Validation is the k-fold Cross-Validation. The k parameter refers to the number of subsets that the input data is split into. Then the result of the model is often summarized as the mean of all the subsets.

The k parameter shall be chosen carefully as a poorly chosen k may result in high bias and high variance. The choice of k is usually 5 or 10 as these values have been shown empirically to yield test error rate estimates that suffer neither from excessively high bias nor from very high variance [16].

K-fold Cross-validation works through shuffling the dataset randomly, then dividing the shuffled dataset into k folds. Then, for all the k-folds the data is trained on the k-1 complementary folds and evaluated on the last fold.

A problem that might occur when utilizing this method is that new data might be qualitatively different from the data the model was trained on. In this study, the dataset consists of seven different subjects for which a 10-fold-cross validation and a 7-fold-cross validation were used. When using 10-fold-cross validation, all of the k-1 folds include data from all the seven subjects, and thus not reflecting a real world example. The effect of including some epochs from the subject expected to predict will result in a better performance than it would be realistic for predicting a whole new and unseen subject data. Thus, to reflect real world performance, the performance metric chosen was a 7-fold Cross-Validation where one fold is one subject.

### C. Architecture of the Scoring Methodology

In this work we used a supervised approach to Machine Learning to solve the problem of automatic sleep scoring. The model was trained on the labeled data by the sleep experts The proposed classifier model is an ensemble between extremely randomized trees and MiniRocket classifiers. The extremely randomized trees model is presented first together with the steps implemented to improve its performance, then the MiniRocket model with its respective improvements. The overall architecture of the methodology implemented in this work is illustrated in figure 2, where the input is the raw EEG data and the output the predicted classes.

**Fig. 2.**
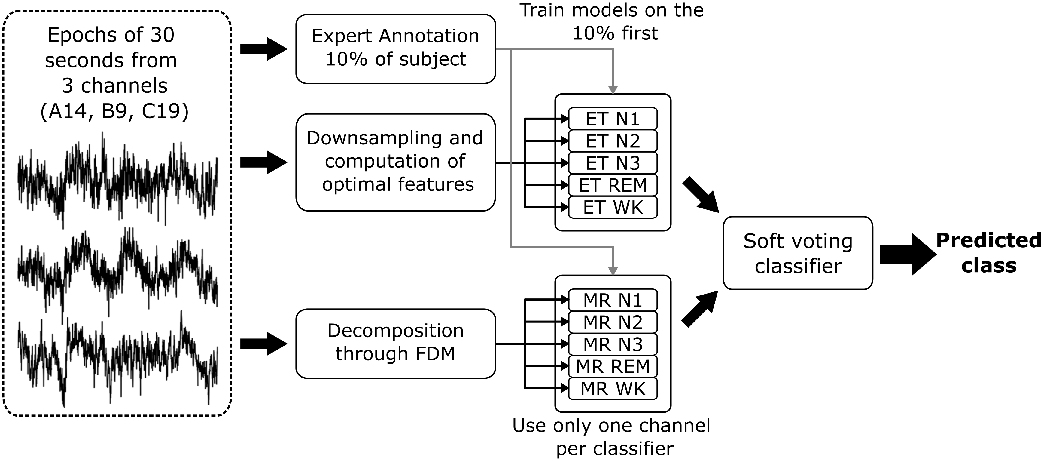
Architecture of the Ensemble Machine Learning Methodology proposed.

#### 1) Extremely Randomized Trees

The Extremely randomized tree algorithm was chosen as it has been successfully used for sleep stage classification and testing proved this classifier to perform better than other classifiers like SVMs and Nearest Neighbors [17].

A few parameters can be tuned to optimize the performance of this classifier. They are the number of estimators, or the number of trees in the forest, and the function to measure the quality of the split. The number of estimators were gradually increased until the accuracy flattened out. 250 estimators were found to be an optimal point as more estimators would increase the model complexity. To tune these parameters the Gini impurity algorithm was used.

Sleep stages appear often in specific sequences after another and this dependence between subsequent epochs can be used to provide extra information to the model. This principle has been implemented through LSTM-layers in deep learning models. A previous study [17], time-shifted the hand-extracted features one step forward and one step backwards, which resulted in a 3-4 percent increase in accuracy in that study. This time-shifting was utilized for the N3 and REM classes.

#### 2) Feature Extraction

The first step for building the model with this classifier was the feature extraction. Feature extraction techniques can be linear and non-linear and in turn be in the time domain, frequency domain and a hybrid of both. A variety of features are reported in the literature for sleep stage classification [18]. The approach in this paper consists of extracting a large number of features and then selecting the features that give the highest accuracy/F1 per class and at the same time reducing the number of features to reduce dimensionality of the data to make it more computationally accessible. To reduce the number of features, an optimization technique based on the NSGA-II algorithm was used [19]. This technique makes possible to represent the data in a reduced dimensional space retaining almost the same information and resulting in a enhanced performance of the classifier.

Figure 2 illustrates the sequence of the step-by-step procedures the input data passes through until optimal EEG channels for highest accuracy are identified.

Regarding the processing of the data used to extract features, the classification model is trained on 1) one feature at a time and 2) using the NSGA-II algorithm on all the extracted features to identify an optimum. The approach in comparing the different individual features is conducted by using the 7-fold Cross-Validation where one fold is one subject. When comparing all the features at once a more standard 10-fold Cross-Validation is used as this is simpler to implement. Figure 3 shows a heat map of the F1 score across all five classes when features were evaluated one at a time.

**Fig. 3.**
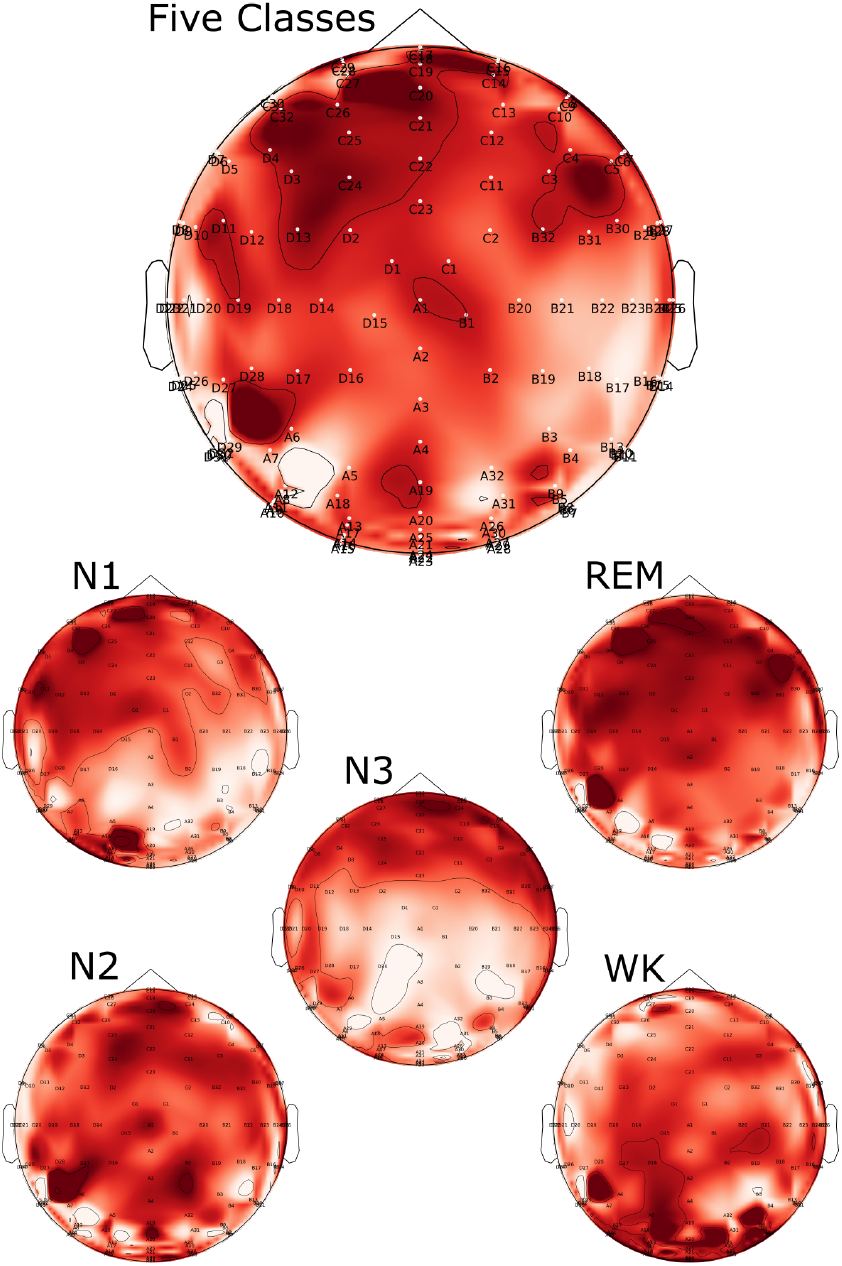
Heatmap of F1 score across all five classes and per each class.

**Fig. 4.**
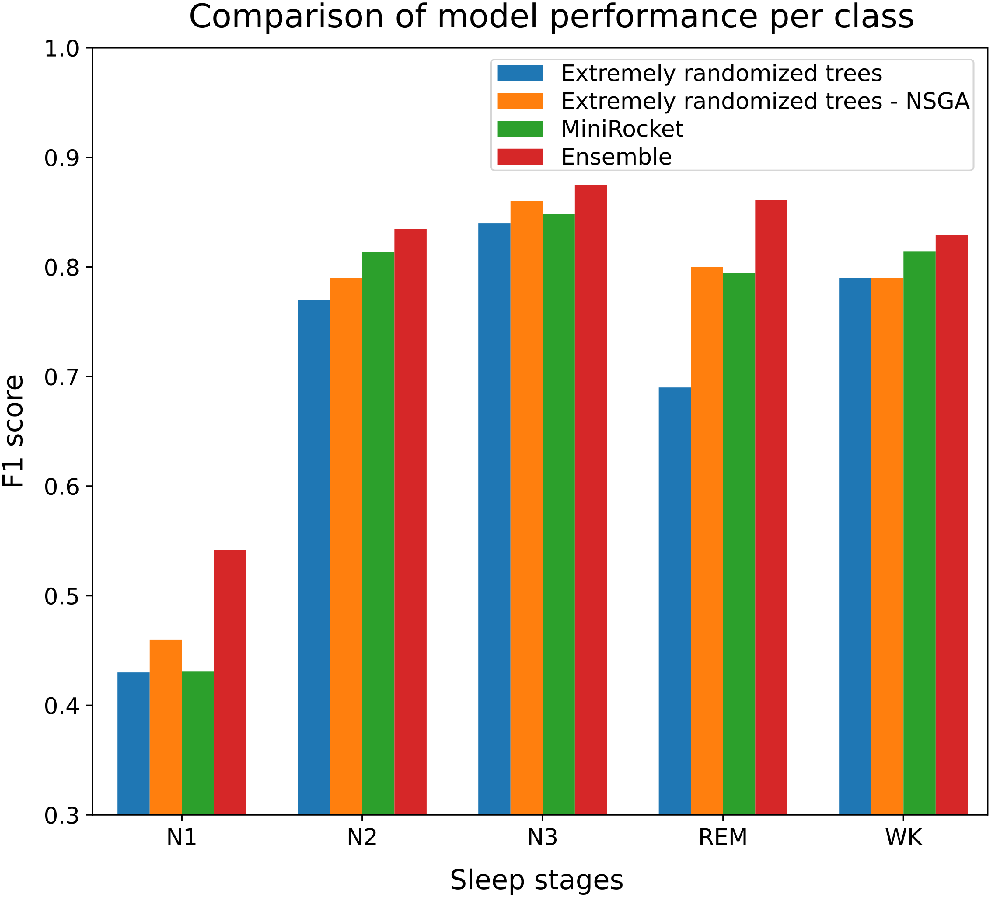
F1 scores of each class for MiniRokcet model, Extra Trees model and ensemble of the two. NSGA algorithm is implemented in the Extra Tree model

#### 3) Resampling

PSG data are unbalanced with respect to the 5 sleep stages, with fewer epochs for stage W and N1 then for N2, N3 and REM. Resampling is a technique that can be used to obtain a more balanced set of data [20]. For that, the raw EEG data sampled at 1024 Hz was resampled to different frequencies to observe any performance difference. As seen in table I the features extracted at 500Hz perform better for almost all features compared to the features extracted at 100Hz, apart from the Petrosian fractal dimension and the permutation entropy where the accuracy are several percentage points higher for the 100 Hz signal. Another aspect when comparing the different frequencies is the computational cost, extracting features on a 500Hz signal takes much longer. This becomes most apparent when calculating the largest Lyapunov exponent and the correlation dimension D2. They were deemed unfeasible for the 500 Hz signal as it took more than a day to extract these on a 28 core CPU. Thus, these two features were only extracted at 100Hz.

**TABLE I.** Comparison of normalization and standardization of the extracted features. the highest score obtained is colored with blue.

| Features | Not processed | Normalized | Zscored |
| --- | --- | --- | --- |
| 75th percentile | 0.561 | 0.429 | 0.441 |
| Analytic entropy | 0.418 | 0.398 | 0.402 |
| Approximate entropy | 0.413 | 0.422 | 0.417 |
| Average spectral power | 0.753 | 0.733 | 0.721 |
| Detrended fractal dimension | 0.451 | 0.434 | 0.429 |
| Higuchi fractal dimension | 0.557 | 0.611 | 0.581 |
| Hjorth complexity | 0.419 | 0.405 | 0.408 |
| Hjorth mobility | 0.424 | 0.421 | 0.387 |
| Katz fractal dimension | 0.411 | 0.400 | 0.405 |
| Kozachenko entropy | 0.405 | 0.403 | 0.401 |
| Kurtosis | 0.405 | 0.404 | 0.382 |
| Log of 8th moment | 0.419 | 0.401 | 0.396 |
| Log of 9th moment | 0.417 | 0.398 | 0.402 |
| Mean absolute deviation | 0.420 | 0.406 | 0.388 |
| Mean | 0.557 | 0.429 | 0.441 |
| Mean of spectral bands | 0.357 | 0.468 | 0.439 |
| Permutation entropy | 0.449 | 0.519 | 0.498 |
| Petrosian fractal dimension | 0.448 | 0.509 | 0.500 |
| Ratio spectral power | 0.688 | 0.705 | 0.706 |
| Relative spectral power | 0.728 | 0.715 | 0.616 |
| Sample entropy | 0.434 | 0.418 | 0.406 |
| Sevcik fractal dimension | 0.430 | 0.413 | 0.416 |
| Skew | 0.391 | 0.375 | 0.357 |
| Slope of the log-log PSD | 0.603 | 0.669 | 0.656 |
| Spectral entropy | 0.481 | 0.444 | 0.434 |
| SVD entropy | 0.426 | 0.418 | 0.417 |
| Variance | 0.326 | 0.394 | 0.401 |
| Number of zero crossings | 0.467 | 0.441 | 0.436 |

#### 4) Normalization

Standardization and normalization are approaches often used as data preprocessing. Standardization is a process where the values are transformed such that the mean of the values is 0 and the standard deviation is 1. The features were also normalized per subject, using the Scaled Robust Sigmoid (SRS) [21], to make the model more general and robust to outliers and to handle interpersonal differences. SRS is a nonlinear transformation that uses median and interquartile ranges instead of the mean and standard deviation. SR is robust against the influence of outliers, and it scales the transformed data into a range between 0 and 1. This was done to make the model more general and robust to outliers and to handle interpersonal differences.

To choose which level of preprocessing and resampling is best for accuracy purposes, again the NSGA algorithm was utilized. The performance measure used was the 10-fold Cross-Validation on the full 5 class classification problem. The results can be seen in table II.

**TABLE II.** Comparison between different levels of preprocessing

| Level of preprocessing | 100 Hz | 100 Hz normalized | 128 Hz filtered | 128 Hz filtered normalized | 500 Hz | 500 Hz normalized |
| --- | --- | --- | --- | --- | --- | --- |
| F1 score | 0.831 | 0.817 | 0.802 | 0.801 | 0.842 | 0.817 |
| Number of features | 15 | 20 | 31 | 42 | 21 | 42 |

Close inspection of these heat maps indicate that some parts of the brain are better suited to extract discriminating information than others. The differences are not extreme, but substantial enough to notice.

### D. MiniRocket

After experimenting with different SoTA models, MiniRocket [22] was found to be the best in terms of performance and runtime. When tuning the MiniRocket hyperparameters and architecture, the number of convolutional kernels was set to 10000. Both 5000 and 15000 convolutional kernels were tested, 5000 decreased the performance with around 2 percent for both the accuracy and F1 score while 15000 increased performance with around 1 percent for both the accuracy and F1 score, but almost doubled the runtime. Thus, for a good accuracy/runtime trade-off 10000 kernels were chosen, or 9996 specifically as it has to be a multiple of 84.

To achieve fast learning of the MiniRocket models Fastai’s implementation of Leslie Smith’s first cycle policy was utilized [23]. Training a deep neural network is a very difficult global optimization problem, where choosing the correct learning rate is crucial for the performance. If the learning rate is low, then training will take a long time and if the learning rate is high it can hinder convergence. In addition, the learning rate is rarely static, it often starts with a higher learning rate to speed up the training and then gradually goes down so that an optimum can be found. Adaptive learning rate methods like ADAM and learning rate schedulers like exponential decay are ways to achieve this. But both those strategies come with a trade-off. Adaptive learning is computationally expensive and learning rate schedulers are set before training starts and thus they will not be able to adapt to the particular problem. Cyclical learning rates combats both these problems by oscillating between reasonable minimum and maximum bounds. A version of a cyclical learning rate is the 1cycle policy where there is just one cycle that alternates between two learning rate steps, and for each cycle the learning rate decreases even further for the next epochs, several orders lower than its initial value [24].

The loss function plays a relevant role in assessing the performance of a model. Sleep scoring is a classification problem that, like many others, must deal with an imbalanced distribution of classes. There are several approaches to deal with a class imbalance. Earlier mentioned is the oversampling of minority classes. This approach is not applicable for MiniRocket as this takes the time series directly as input. Methods for creating an artificial sample of a time series exist, but was deemed outside the scope of this study. Undersampling is also an option, but with the already limited dataset, this was not tried. The use of the Focal Loss objective function was proposed in [25] to deal with class imbalance. This is based on giving more weight to all the hard (false negatives) samples than the easier ones(true negatives). The degree to which this is done is decided by the gamma hyperparameter. The alpha parameter is a weighing factor per class, this is usually set by the inverse class frequency. The Focal Loss was combined with Dice Loss, Dice Loss to combat class imbalance. During testing, the class weighting factor was proven to be agressive, giving the smaller classes a very high recall, but a low precision score. Thus, all the weights were increased by 1 such that the relative difference would be smaller. For the 5 class classification problem introducing the new Loss function performed equally. However, for the one vs all strategy (reference) introducing a new loss function improved the performance by a lot for some of the classes of some of subjects. For subject 6 classifying REM against the other classes using Cross Entropy Loss resulted in an F1-score of 0.76, while when using the new loss function an F1 score of 0.85 was achieved.

#### 1) Preprocessing

For the input to this model the data was first decomposed in sub-bands using the Fourier Decomposition Method (FDM) [26]. The following frequency bands were extracted: [30-48] Hz, [12-30] Hz, [8-12] Hz, [4-8] Hz, [0,4] Hz, [0.5-2] Hz, [2-6] Hz, [12-14] Hz. These bands were selected based on the AASM manual to represent the delta, theta, alpha and beta waves. Additional bands were added, 0.5-2 Hz for slow wave activity, 30-48 Hz for differentiating the wake stage based on [27], 2-6 Hz to detect sawtooth waves and 12-14 Hz to detect sleep spindles.

The FDM improved the accuracy for MiniRocket by a substantial margin. On the raw signal from the A28 channel the model achieved an accuracy of 0.686930 and an F1 score of 0.686317. After decomposition with FDM signal from the A28 channel achieved an accuracy of 0.806560 and an F1 score of 0.799751. Thus, using the FDM was the chosen method.

#### 2) Channel Selection

As opposed to the Extremely randomized trees model, the MiniRocket model did not improve by a significant margin when using more channels. Among the three channels available, the binary classifiers for the N2, N3 and REM classes are trained on data extracted from the C19 channel. While the N1 classifier was trained on the A14 channel and the WK class was trained on the B9 channel. This choice is motivated on the F1-score of the Extremely randomized trees models which were trained on one and one channel for all the five classes and the N1 binary classification model got the highest score on the A14 channel and the WK binary classification model got the highest score on the B9 channel. The rest of the classes performed best on the C19 channel.

### E. Channel and Feature Optimization

In this work, the entire space of 128 EEG channel positions was used as search space for minimizing the information required to obtain high accuracy classification. Most studies that report reduced number of channels start from the reduced subset of the given PSG channel configuration used for recording. Here the recordings were done with 128 EEG channels, and the NSGA-II optimized search, the goal is to achieve the highest accuracy/F1 score while at the same time using the minimum number of EEG channels. This approach of minimizing the number of electrodes has proven to be effective to identify subset of channels while retaining high accuracy in multiple problems [28], [29]. Using the NSGA algorithm, the number of channels were constrained around 5,4,3,2,1. The F1 was determined by using the extremely randomized trees algorithm with a reduced number of features (10), extracted from the 100Hz resampled normalized signal. The features included, mean, average power of the EEG bands (5), Petrosian fractal dimension, permutation entropy, analytic entropy and Higuchi fractal dimension. These features were chosen as they yielded a high accuracy/f1 score on their own.

In parallel to the optimized search of channels by the NSGA algorithm, every channel was also individually used to calculate the accuracy/F1 per class per channel. This was done to have an oversight over the best channels per class. The result of this is displayed in the heatmap of 3, where all the values are normalized through the use of the Scaled Robust Sigmoid to accentuate the differences. The results for the best and worst performing channels are displayed in Table ref.

Then, to select the best features per class, the NSGA algorithm was again used. For each class, the objective function aimed to minimize the number of features and to maximize the F1 score.

The algorithm returned the optimal features after 350 generations The total number of features were 145, 48 per channels and the spindle feature

## III. Results

### A. Classification Performance

As indicated in section II-B, to measure the quality of the automatic scoring, the F1 score was found to be the most suitable, as it represents a balance between recall and precision. Figure 3 illustrates the F1 computed over all 5 classes on the validation/testing of the dataset for all stages of sleep. All stages show high performance except for N1. We consider this to be within reasonable levels considering the low human interscorer agreement for N1 [30].

A technique that was tried was to train the model on a portion of the unknown subject to see if the different subjects had different characteristics and those characteristics would increase the overall performance of the model. This was tried because the F1 score obtained through the 10-fold Cross-Validation was not repeated with the 7-fold Cross-Validation, and the difference is that the 10-fold Cross-Validation contains epochs from all subjects. With the 10-fold Cross-Validation the F1 score was 0.84 and for the 7-fold Cross-Validation the F1 score was 0.79. This suggest that training on a portion of the unknown subject can increase the performance by several percentage points.

### B. Results of the NSGA Optimization

When using five channels the optimal F1 score was 0.826, when using four channels the F1 score was 0.826, when using three channels the F1 score was 0.821, when using two channels the F1 score was 0.810 and when using only one channel the F1 score was 0.782. This is presented in figure 6.

**Fig. 5.**
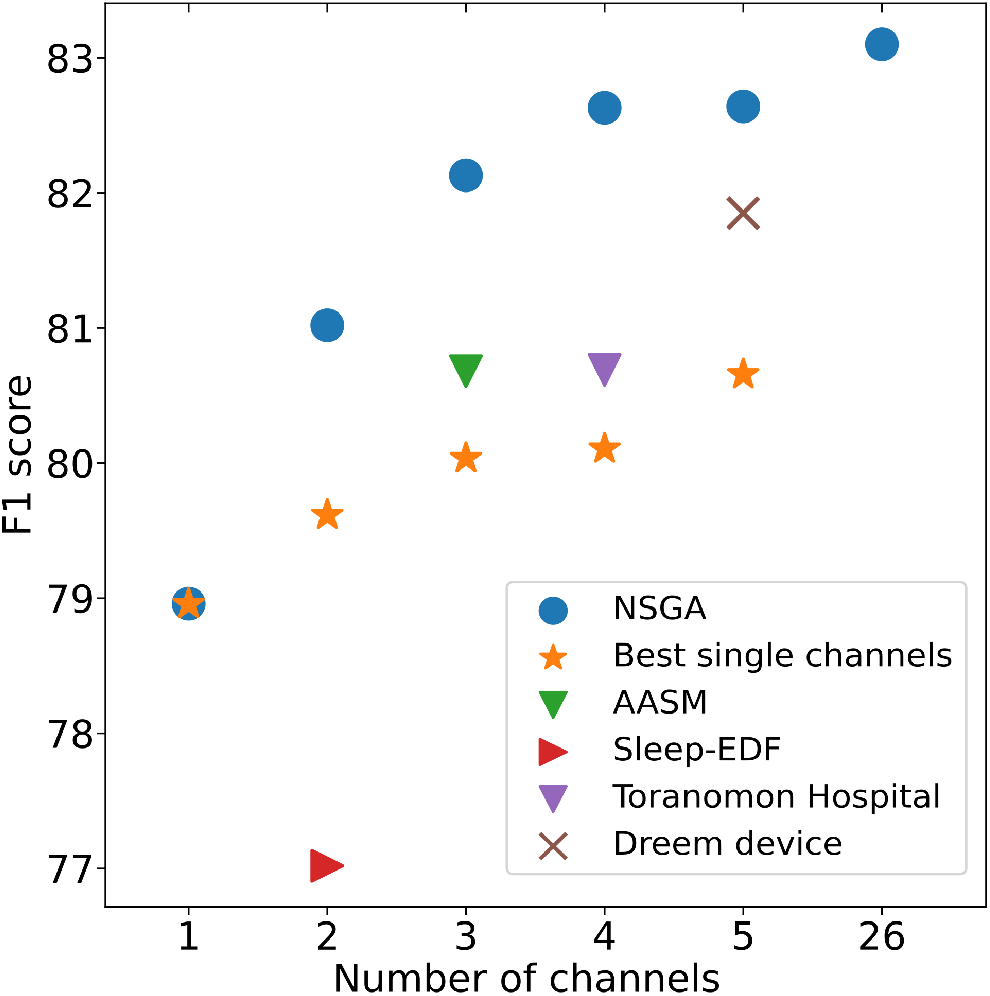
F1 scores of the 5 best EEG channels resulting from the NSGA algorithm optimization (blue circles) compared to best 1-5 channels when individually evaluated from F1 Heat map (yellow stars), Channels recommended by AASM (green inverter triangle), channels used in the Slee-EDF dataset (red triangles) and Channels used for sleep scoring by Toranomon Hospital (inverter violet triangle).

**Fig. 6.**
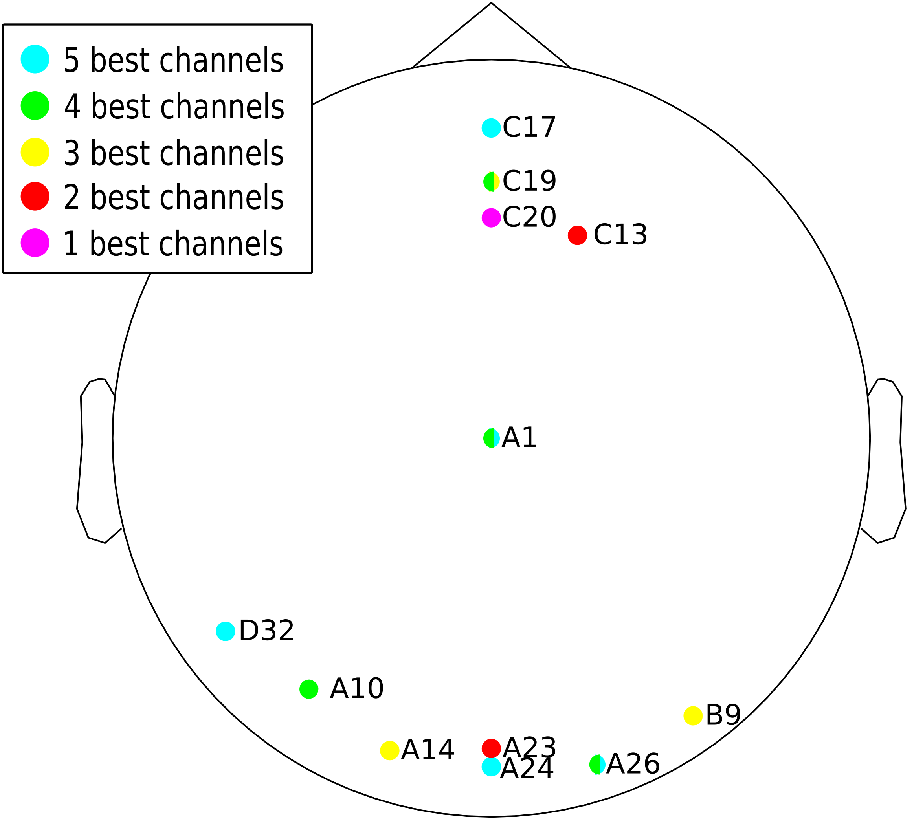
Channel locations of the best 5, 4, 3, 2 and 1 channels.

Since the objective of this study is to use a minimal set of EEG channels three channels were chosen as the difference from two channels is larger than the difference between five channels.

## IV. Discussion

The results of channel selection using the NSGA optimization algorithm shed some light into the impact of the number of EEG channels on the classification performance. Not only the number of EEG channels but their location on the scalp play a role in accuracy and this role depends on the selected features and on the classification technique used. The result is highly dependent on the parameters associated with the methodology used (certain combination of features will work better with certain channels), the solution is not unique and is highly parameterized. Reasonably good F1 scores can be obtained for different channel combinations (50, 20, 10, 5, etc) by constraining the solution space around those channel numbers. These solutions represent local minima that all offer similar values of F1 score which can be suitable for sleep scoring. In this work, and with the idea to identify a minimum set of optimized channels that can facilitate adoption in the clinics, the optimization was constrained around 5 channels with F1 scores as illustrated in figure 5. Several previous studies report the use of different EEG channels with or without EOG and EMG channels. Most of these works focus on EEG single-channel with different single channel proposals [8]. This review paper shows that the classification performance generally increases with a multi-channel approach but not very significantly and the results of this work confirms the same. The work presented here is the first attempt to systematically search for the optimal single and multi-channel subsets for a low number of channels combinations. Figure 5 plots those solutions together with multi and single-channels reported in the literature (using different classification methodologies). It is evident that when those reported channels are evaluated with our proposed methodology, the accuracy is inferior (for the same number of channels), which reveals the strong dependency of the solution on the specific parameters of the used classification method. In the same plot it can also be observed that, as the number of NSGA channels increases (4 to 5), there is only an incremental improvement of F1 score (the performance appears to plateau). However, the improvement of F1 score is more salient when channels increase from 1 to 2, and 2 to 3, lowering down from 3 to 4 channels. Although many studies have shown that multi-channel EEG with EMG and EOG leads to an increase of performance, the improvement is often marginal and adding one more channel may compromise the computational cost without leading to a better performance. In clinical settings and for home-based sleep devices, solutions with fewer channels will be preferred and an important question still under debate is ”which fewer channels to use?”. From previous works and ours, we can say that there is no ”one-size-fit-all” solution and that extensive training on larger datasets with high heterogeneity (including subjects with sleep disorders, local sleep, etc) should be considered before deciding on a given multi-channel combination. For example, a single-best channel or even a few number of best channels might not be suitable for scoring scenarios of local sleep or for scoring sleep on subjects with sleep disorders [31], [32].

In this context, the flexibility offered by the NSGA algorithm may be further exploited to identify more general models and best suited channels for various sleep scenarios. One way of doing this is by optimizing the parameters of the ML learning models, the length of epoch used, and even the classifiers to be used for the given scenario, by allocating each of them to a single chromosomes of the NSGA.

The optimal features found by utilizing the NSGA algorithm were slightly unexpected when compared to the results found in previous studies. However, most of the features listed in those studies were good at discriminating one class from another and not one class from all the other classes which is the case for sleep stage classification. The features listed were also often described at discriminating sleep stages from one another on their own, and the results found in this study could indicate that a combination of different features outperform a single feature in discriminating between different sleep stages. These combinations can be uncovered though the use of the NSGA algorithm.

The combination of feature-classifier-epoch size-data set characteristics will significantly impact the classification accuracy. Possible explorations to extend the model capabilities include unsupervised classification approaches similar to the ones presented in [13], [27], [33]. For these, extensive training using heterogeneous data-sets will be necessary.

This study shows that automatic sleep scoring is able to to reach a SoTA performance. Nevertheless, all these approaches, both extremely randomized trees and MiniRocket classifier, reach a similar performance level and one should question if it is possible to reach a higher level, based on the inter-rating consistency [34]. This could mean that the labels can be inconsistent and that the scoring standard is not clear enough. The problem with the scoring standard had been addressed in the study of [13] and should be explored further. And a true unsupervised data-driven approach should be further explored.

As the field of machine learning is continuously expanding and improving there might come new models along which will give a consistently high score, so this should also be explored with new and improved machine learning models. Also as mentioned the use of different loss functions might optimize the performance of the SotA MiniRocket classifier.

Another approach could be to utilize architectures like DOSED [35], to detect micro-architecture events with a high accuracy and thus maybe score sleep through a flow chart following the existing rules set by the AASM.

## V. Conclusion

We demonstrated in this work that it is possible to achieve reasonably good scoring accuracy of sleep stages with an optimized minimum set of EEG channels identified by an optimization procedure. The ensemble model, compared to its single components independently, provided a quality of scoring comparable to that of human experts. The optimization routine offers a systematic way to select only EEG channels that contain the most relevant information for accuracy.

The results obtained in this study are comparable to results obtained by other studies, but the analysis of which channels perform best is previously unseen and should be delved into deeper.

Since the dataset of this study has a uniqueness when it comes to the number of channels available and tsuhe sampling frequency and these aspects of the dataset were utilized to maximize the performance, the results are not directly comparable to the results obtained using other datasets. Although the performance metrics were similar to the ones obtained in the other studies, what this study has shown is that some techniques utilized here that increase performance can be applied to other sleep stage classification studies, like the ensemble of classifiers that complement each other, the inclusion of oversampling techniques or using different features per binary classifier which is in turn combined through an all against one approach.

However, based on the results of other automatic sleep stage classification models and the inconsistency of human scoring, some weakness of the established scoring rules might have been uncovered. This could call for the implementation of an unsupervised data-driven approach, which already has some traction in the sleep study field.

With the added value of an optimization routine that extended the search space to the entire 128 channels to identify optimal combinations of channel number and features, a road is open for a more systematic search of fewer channels that can facilitate on-line implementations and lead to the design of new and reliable home-based sleep devices and to the adoption of sleep devices in the clinics.

## Acknowledgment

AS was supported by Enabling Technologies - Norwegian University of Science and Technology, under the project ”David versus Goliath: single-channel EEG unravels its power through adaptive signal analysis - FlexEEG”.

## Footnotes

1 https://biosemi.com/pics/cap_128_layout_large.jpg

